# Automated image analysis method to detect and quantify fat cell infiltration in hematoxylin and eosin stained human pancreas histology images

**DOI:** 10.1101/2021.12.13.472341

**Authors:** Roshan Ratnakar Naik, Annie Rajan, Nehal Kalita

## Abstract

Fatty infiltration in pancreas leading to steatosis is a major risk factor in pancreas transplantation. Hematoxylin and eosin (H and E) is one of the common histological staining techniques that provides information on the tissue cytoarchitecture. Adipose (fat) cells accumulation in pancreas has been shown to impact beta cell survival, its endocrine function and pancreatic steatosis and can cause non-alcoholic fatty pancreas disease (NAFPD). The current automated tools (E.g. Adiposoft) available for fat analysis are suited for white adipose tissue which is homogeneous and easier to segment unlike heterogeneous tissues such as pancreas where fat cells continue to play critical physiopathological functions. The currently, available pancreas segmentation tool focuses on endocrine islet segmentation based on cell nuclei detection for diagnosis of pancretic cancer. In the current study, we present a fat quantifying tool, Fatquant, which identifies fat cells in heterogeneous H and E tissue sections with reference to diameter of fat cell. Using histological images of pancreas from a publicly available database, we observed an intersection over union of 0.797 to 0.966 for manual versus fatquant based machine analysis.

**Author Summary:** We have developed an automated tool, Fatquant, for identification of fat cells based on its diameter in complex hematoxylin and eosin tissue sections such as pancreas which can aid the pathologist for diagnosis of fatty pancreas and related metabolic conditions. Fatquant is unique as current fat automated tools (adiposoft, adipocount) works well for homogeneous white adipose tissue but not for other tissue samples. The currently available pancreas analysis tool are mostly suited for segmentation of endocrine β-cell based on cell nuclei detection, extracting colour features and cannot estimate fat cell infiltration in pancreas.

**Graphical Abstract:** 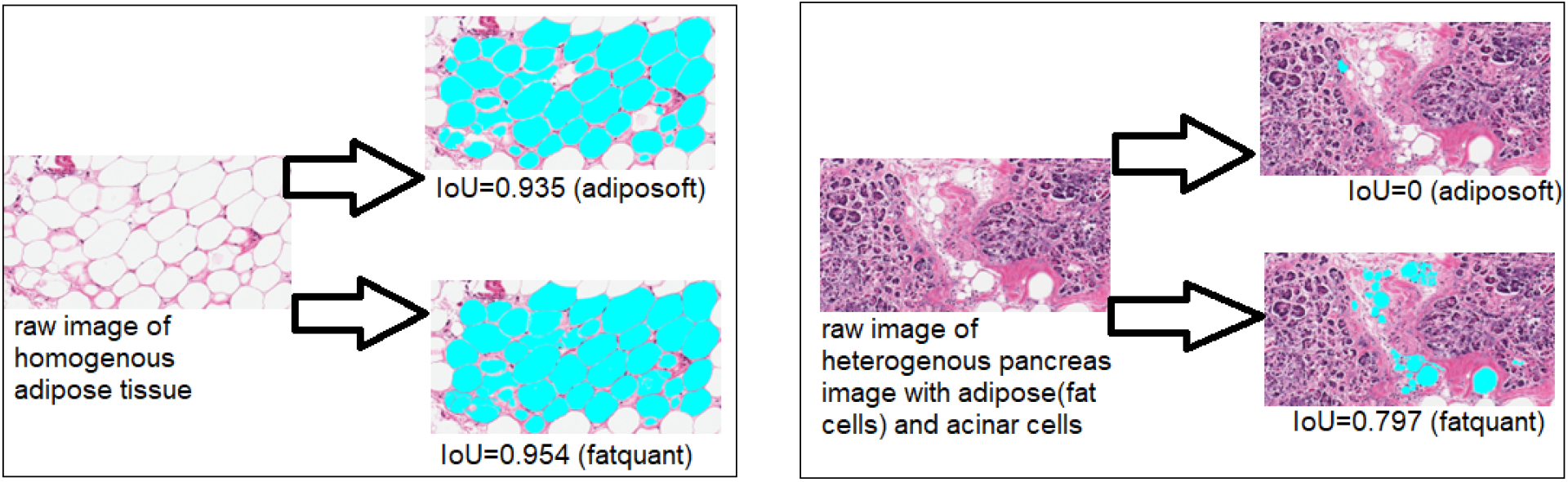

Currently available fat quantification tools like adiposoft can analyze homogenous adipose tissue (left) with intersection over union (IoU) of 0.935 and 0.954 with adiposoft and fatquant, respectively. While in heterogenous tissue (e.g. pancreas on right) which contains adipose (fat cells), acinar cells, adiposoft fails to detect fat cells with IoU=0 while fatquant had IoU=0.797.

## 1. Introduction

The accumulation of fats especially in the abdominal area causes insulin resistance leading to deposition of fats (steatosis) in the pancreas, inflammation and finally fibrosis leading to non-alcoholic fatty pancreatic disease (NAFPD). Accumulation of pancreatic fat may lead to pancreatitis, diabetes mellitus or pancreatic cancer (Paul and Shihaz 2020). Pancreatic steatosis can be diagnosed on ultrasound, computed tomography (CT) scan or magnetic resonance imaging (MRI) but pancreatic biopsy remains best method to detect pancreatic fat concentration (Tariq et al 2016, Paul and Shihaz 2020). The consequence of pancreatic fat infiltration might provoke a decrease in endocrine (β-cell) number and function, leading to more rapid progression to diabetes (Yu and Wang 2017). Sudies suggest NAFPD as an early marker of glucometabolic disturbance (Yu and Wang 2017). Pancreas transplantation is the only way to treat type 1 diabetes (T1D) but fat infiltration in pancreas remains a risk factor that can affect the clinical outcome (Verma and Papalois 2011, Dholakia et al 2017). Fatty pancreas has a prevalence of 35% and may be contributing factor for the malignancy and the metabolic syndrome (Lesmana et al 2015). The histological and MRI tools exhibit good agreement in detecting fat in pancreas (Fukui et al 2019, Virostko 2020) but the latter cannot show fat accumulation at the cellular level and is expensive. Decrease in fat content in liver and pancreas was associated with non-diabetic blood glucose control in people with type 2 diabetes (Taylor et al 2018).

Hematoxylin and eosin (H & E) is a widely used histological tissue staining technique for medical diagnosis and scientific research. Hematoxylin stains cell nuclei blue while eosin stains the cytoplasm and connective tissue pink thus allowing microscopic differentiation of tissue cytoarchitecture in sections. The analysis involves manual examination by pathologist to ascertain presence/absence of disease markers and or grading of disease progression which is semi-quantitative and subjective in nature (Gurcan et al 2009).

To complement the current manual assessment, several digital tools have been developed such as Adiposoft (Galarraga et al 2012), and AdipoCount (Zhi et al 2018). However, they cannot efficiently identify fat cells in complex tissues such as pancreas or liver as white adipose tissue is generally homogeneous (Figure 1) making tissue segmentation relatively easier (Glastonbury et al 2020). Such tool developed based on nuclear displacement and lipid droplet size analysis exists for automated analysis of fat cell infiltration (steatosis) in H and E liver images (Nativ et al 2014). Unlike liver tissue, where fats accumulate in hepatocytes, in pancreas fat infiltration and deposition occurs both in acinar and islet cells (Catanzaro et al 2016). The ratio of accumulated pancreatic fat area relative to exocrine gland (acinar and duct tissue) area was significantly increased in obese mice model (Matsuda et al 2014).

**Figure 1.**
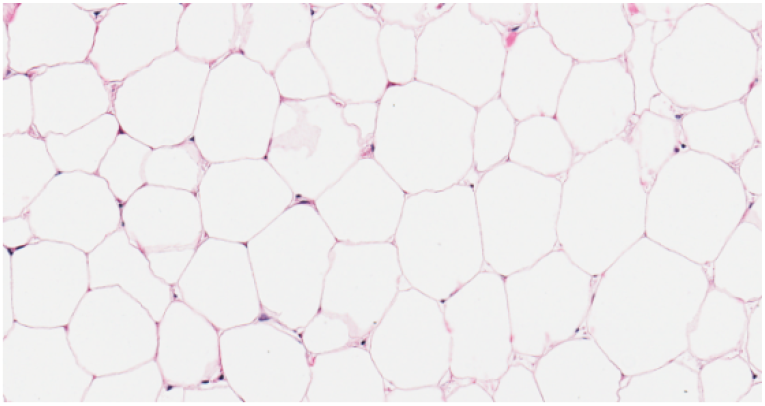
Homogenous adipose tissue

Usually, these algorithms involve splitting the image into various color channels with the red channel binarized using automatic thresholding method to separate the bright pink fat areas from dark purple-bluish cell nuclei. Subsequently, a watershed algorithm is applied to fillup missing fat cell membrane and improves cell count (Galarraga et al 2012, Zhi et al 2018). The output of these processes includes the labels and statistical analysis of individual cells. These tools have been successfully applied to white adipose tissue, however, other organs like liver, pancreas, lungs have been challenging due to heterogeneous cell types (Figure 2).

**Figure 2.**
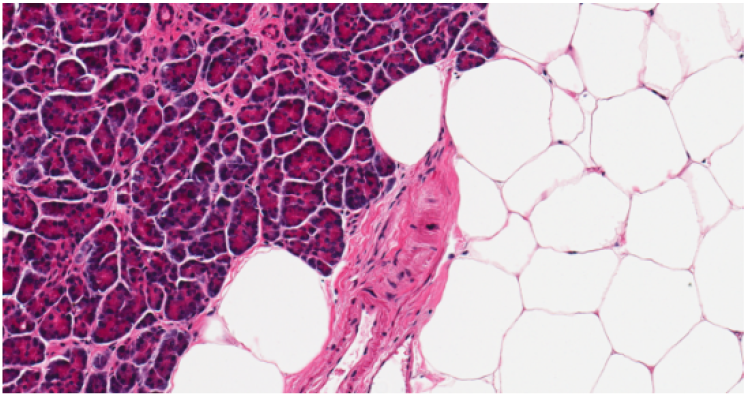
Heterogenous pancreatic tissue with adipose and acinar cells

Pancreas is a heterogenous tissue and manual analysis of regions is a tedious process that lacks reproducibility (Apaolaza et al 2021). The existing methods of pancreatic islet segmentation depends on cell nuclei detection, then a classifier is applied to recognize different cell types but is based on assumption that islets have high density cells (Floros et al 2009, Rechsteiner et al 2014). Recent studies have utilized a supervised learning framework for islet segmentation in H & E stained pancreatic images to partition images into superpixels and extract color-texture features, process them, and finally a linear support vector machine is trained and applied to segment testing images (Huang et al 2016). Moreover, QuPath software allows identification of endocrine islet cells in immunofluorescent images of pancreas tissue (Apaolaza et al 2021), but these tools does not allow quantification of adipose (fat) cells in pancreatic images. In the current study, we have developed an automated tool, Fatquant, in which the fat cells were identified in processed images by calculating the diagonal of a square circumscribed by circle.

## 2. Materials and Methods

The H and E images for the analysis were downloaded from a publicly accessible Genotype-Tissue Expression (GTEx) Portal (Broad Institute, Cambridge, MA, USA) using the Histology Viewer tool (Consortium 2015). The GTEx tissue image library contains high-resolution histology images for various tissue types from several postmortem donors. Spherical or oval white spaces were categorized as fat (adipose) cells while large and irregular white spaces were grouped as artifacts. A sample size of ten pancreas histology images were analyzed from slide IDs GTEX-11DXZ-0826, GTEX-1122O-0726, GTEX-1117F-1726, GTEX-117YW-0926, GTEX-13PVQ-2026, GTEX-13FHP-1926 and GTEX-11WQC-0926. The images have magnification of 20x (0.4942 mpp—microns per pixel) (Badea and Stanescu 2020). The sample images were taken using the Snapshot tool of Aperio ImageScope software. The source code was written in Python version 3.8.0 (Python Software Foundation, Beaverton, Oregon, USA) with Core i7 3rd Generation CPU, 8 GB RAM and Intel HD graphics 4000 GPU.

### 2.1. Data and code availability

All the images, annotations along with relevant data code can be found at the following GitHub repository: https://github.com/anniedhempe/Fatquant. The procedure to run this tool is mentioned in the Readme file.

### 2.2. Image processing

The procedure for image processing is briefly demonstrated in the flow chart shown in Figure 3.

**Figure 3:**
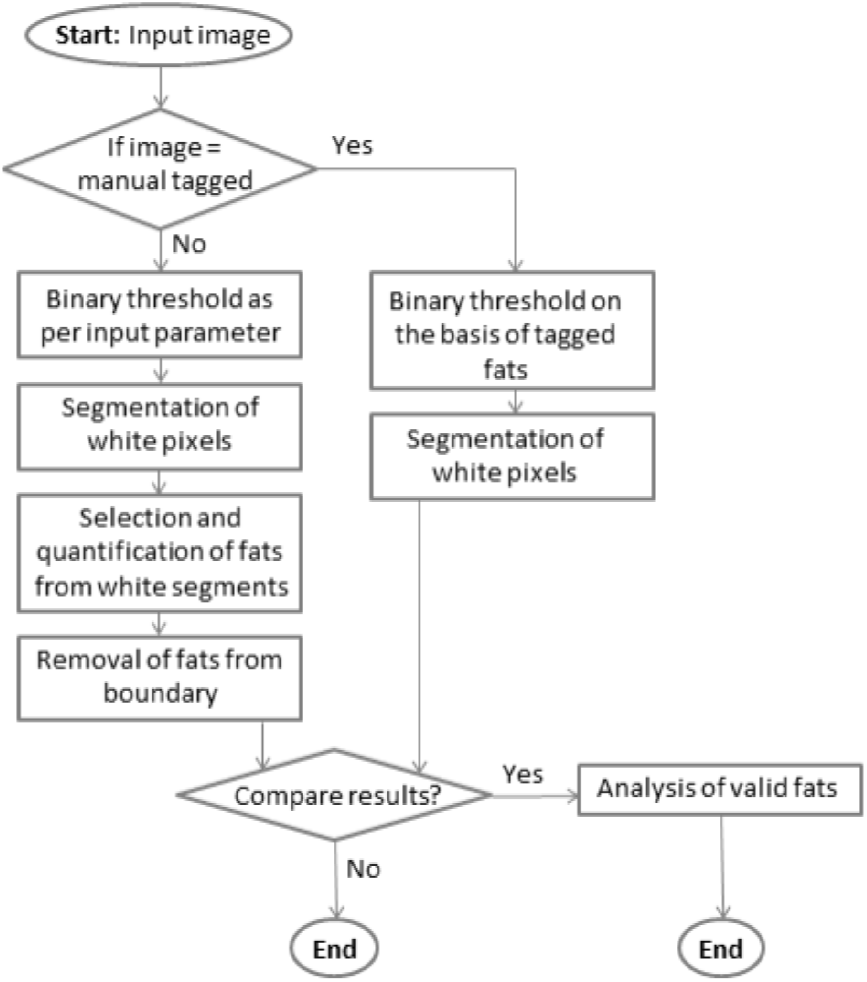
Flowchart depicting the image processing.

The steps used are elaborated below.

Figure 4 is a sample raw image of dimension 1716 × 905 pixels from whole slide GTEX-11DXZ-0826. Analysis on this image is referred while explaining the image processing steps.

**Figure 4.**
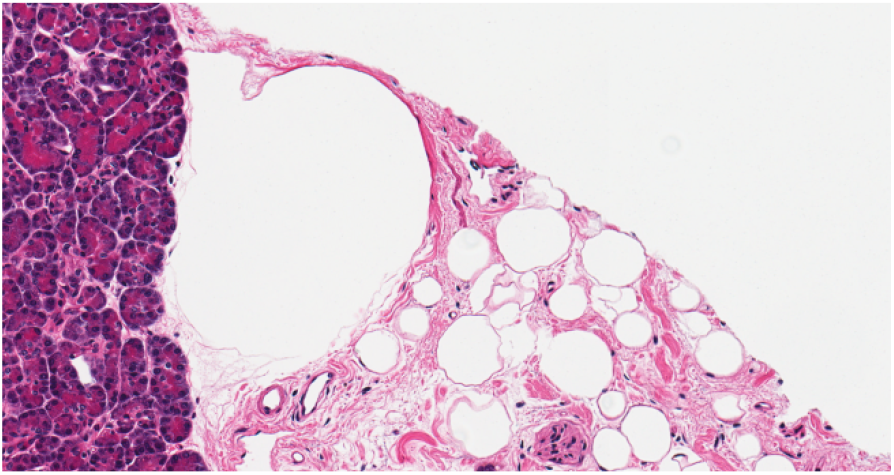

Figure 5 is an altered form of the image in Figure 4 where the valid fat cells are manually tagged with yellow color (RGB: 255, 255, 0).

**Figure 5.**
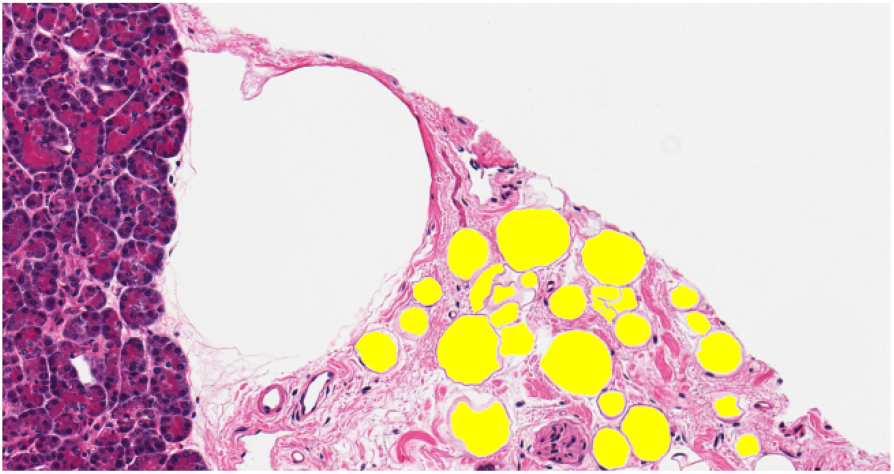

#### 2.2.1. Binary thresholding of input image

The color of fat cells in images from GTEX portal ranges approximately between 225 to 255 grayscale values. There can also be other parts of pancreas which has the same range of color. But applying binary threshold on an image can help in getting rid of many unwanted parts. The pixels of an image whose color values are at least equal to the input parameter value of threshold (e.g. 227) are taken into consideration for further processing and are assigned a new grayscale value 255. The other pixels are assigned value 0. Figure 6 is a thresholded image of Figure 4 with parameter value 230. Figure 7 is also a thresholded image but of Figure 5 where pixels representing the tagged fat cells are assigned grayscale value 255 and the rest is assigned value 0.

**Figure 6.**
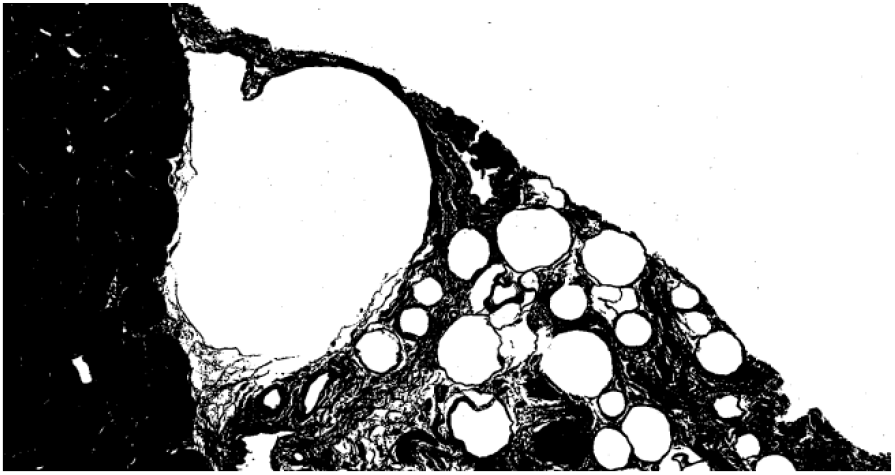

**Figure 7.**
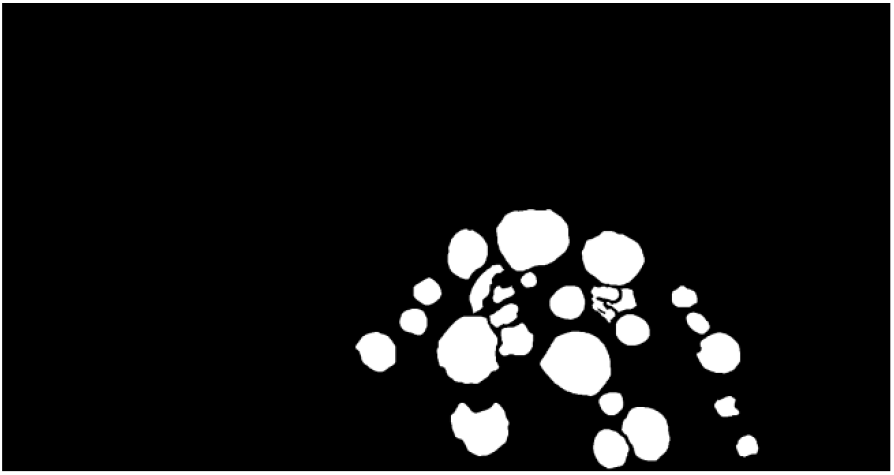

#### 2.2.2. Segmentation of white pixels from thresholded image

White pixels are initially segmented by combining tile rendering with scanline rendering and then identifying possible merge of segments in a tile with their immediate neighbors. Tile rendering has been implemented in this system as it helps in reducing time complexity for segments covering large area.

Processing time for segmentation was tested with four sizes of square tiles, which were of length 35 pixels (processing time: 24.23 seconds), 50 pixels (processing time: 15.32 seconds), 70 pixels (processing time: 12.88 seconds) and 100 pixels (processing time: 15.37 seconds). In this experiment, tile size of 70 pixels was used for analyzing all the images since it took the least amount of time.

After an input image is divided into multiple tiles, scanline rendering is performed within each tile to identify white pixels and form possible segments with their neighbors on left or top. In this experiment, left neighbors are given first preference. Once scanline rendering is performed till the last row of a tile then the identified segments are merged on the basis of their vertical neighbors. Segments which have only diagonal neighbors with another segment and not vertical, are not merged.

Figure 8 is a diagrammatic representation of segmentation performed on a square tile of length 15 pixels. Figure 8 (a) represents a thresholded image where segmentation is to be performed. Figure 8 (b) shows six segments created while iterating once till the last row. Figure 8 (c) shows total segments getting reduced to four due to merging.

**Figure 8 (a).**
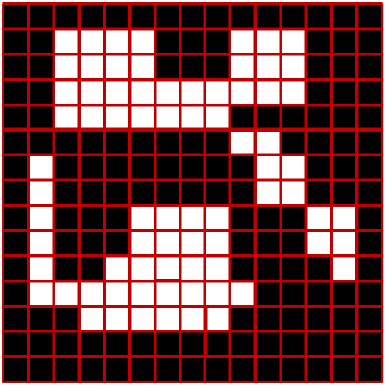

**Figure 8 (b).**
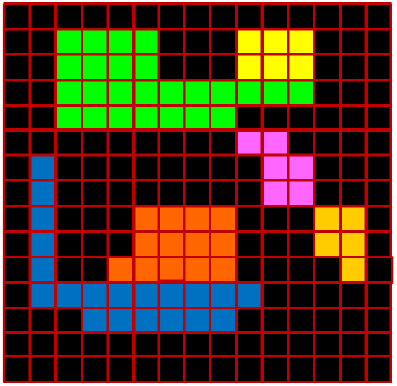

**Figure 8 (c).**
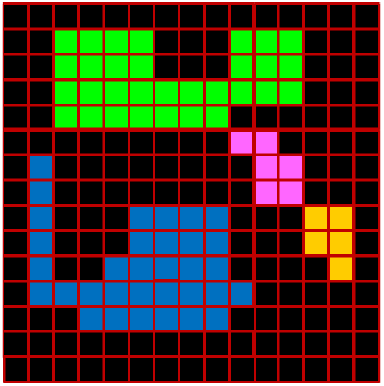

Figure 9 shows segments within tiles and the magnified part is one tile. Figure 10 shows fully merged segments of the thresholded image and varying colors (non-black) in it represent different segments.

**Figure 9.**
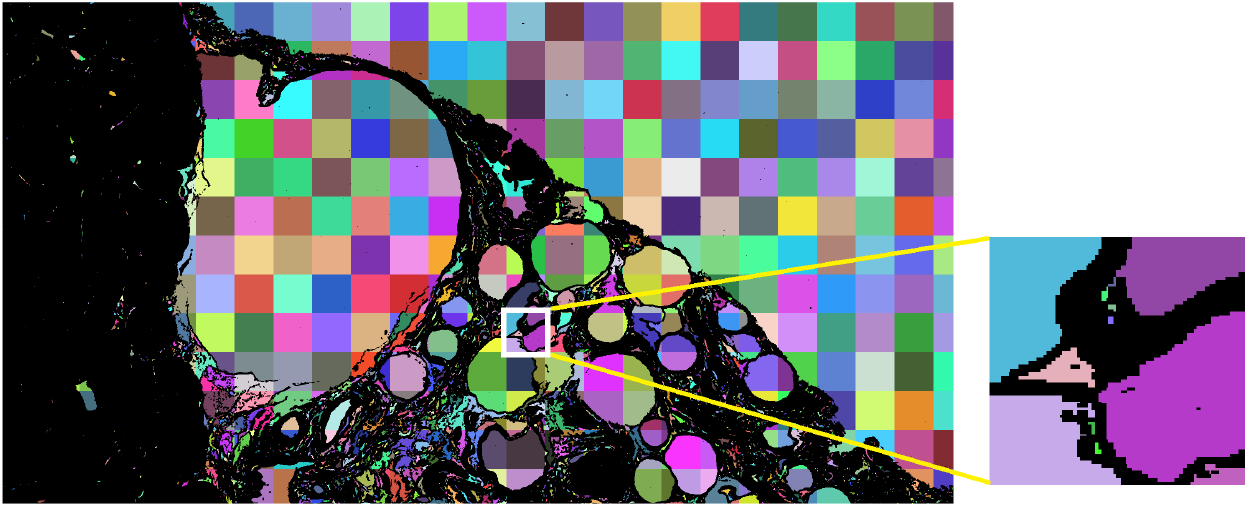

**Figure 10.**
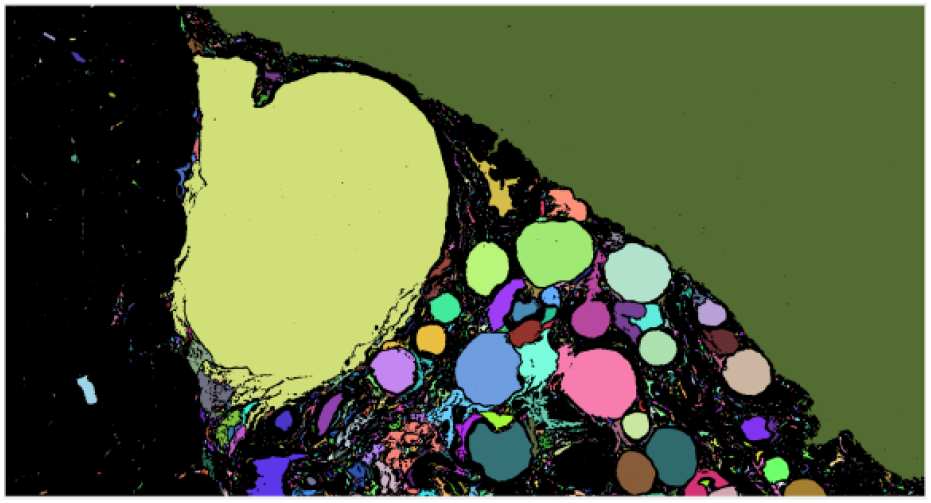

#### 2.2.3. Selection and quantification of fats from white segments

The system uses image kernel to select probable valid segments. But before applying kernel, the system determines the smallest dimension that encompasses all the segments. The kernel’s position is only updated by one pixel (horizontally or vertically) in each iteration so determining the smallest dimension for traversal reduces the time complexity. This dimension is determined by identifying positions of first white pixels in thresholded image from every direction (i.e. top, bottom left and right). These four positions denote maximum coverage of segments in each direction. Figure 11 is a diagrammatic representation of the method discussed where the Red colored rectangle represents smallest dimension.

**Figure 11.**
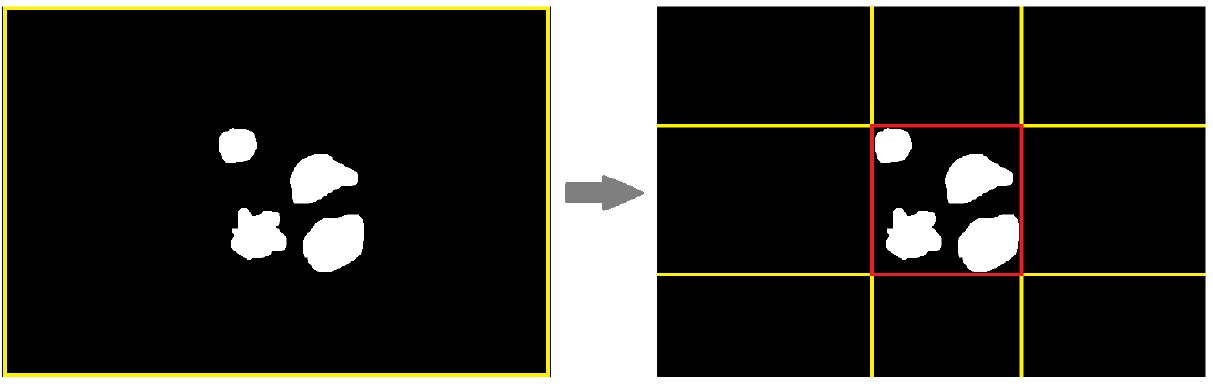

The system uses a square shaped kernel which is supposed to fit inside the boundaries of valid segments. The side length of that square is determined by input fat diameter values (minimum or maximum) as these diameters equates to diagonal of that square. Side length of a square, can be calculated as:

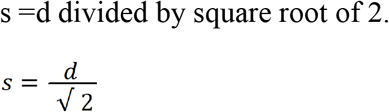

where s = square side length; d = diameter. This square can be assumed as the largest square that can get inscribed in a circle of given diameter. An elliptical kernel may be a more precise choice for identifying fats but since square kernel is easier to handle so it has been chosen.

The system refers to a minimum diameter value to select segments where a square kernel having side length as per this diameter can fit somewhere in their regions. Then the system refers to a maximum diameter value to discard segments from the selected list where a kernel having side length as per this diameter can fit somewhere in their regions. This means a segment which has narrow areas in many of its portions, but has very wide area in one of its portion can also get discarded if a square kernel as per the maximum diameter can fit in that portion. Figure 12 is a diagrammatic representation of this process performed on an image of dimension 15 × 15 pixels. Figure 12 (a) has seven segments with White colored pixels out of which valid segments are to be selected. Figure 12 (b) has three segments marked with Cyan color which denotes segments getting selected as per minimum diameter. The kernel size is of length 3 pixels. So, four segments do not get selected as the square kernel fails to fit inside the boundary of any of these segments. Figure 12 (c) has only two segments marked with Cyan color which denotes selected segment getting discarded as per maximum diameter. Here the kernel size is of length 4 pixels. So, a segment which can fit a kernel of length greater than 4 pixels is to be discarded. The previously selected segment which gets discarded could fit a kernel of length 5 pixels. Figures 13 (a) and (b) are outputs generated from the sample image with minimum (27 pixels) and maximum (130 pixels) diameter respectively. The identified fats are marked with Cyan color (RGB: 0, 255, 255).

**Figure 12 (a).**
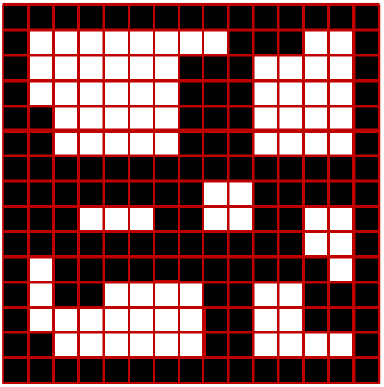

**Figure 12 (b).**
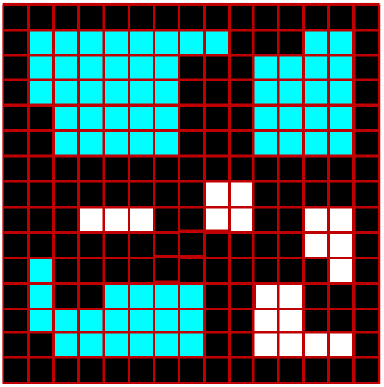

**Figure 12 (c).**
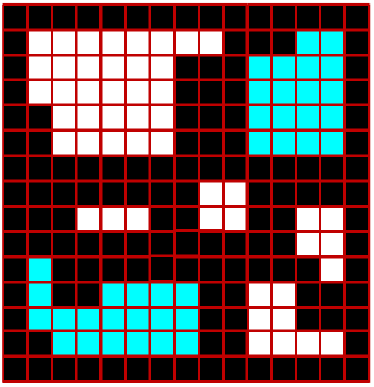

**Figure 13 (a).**
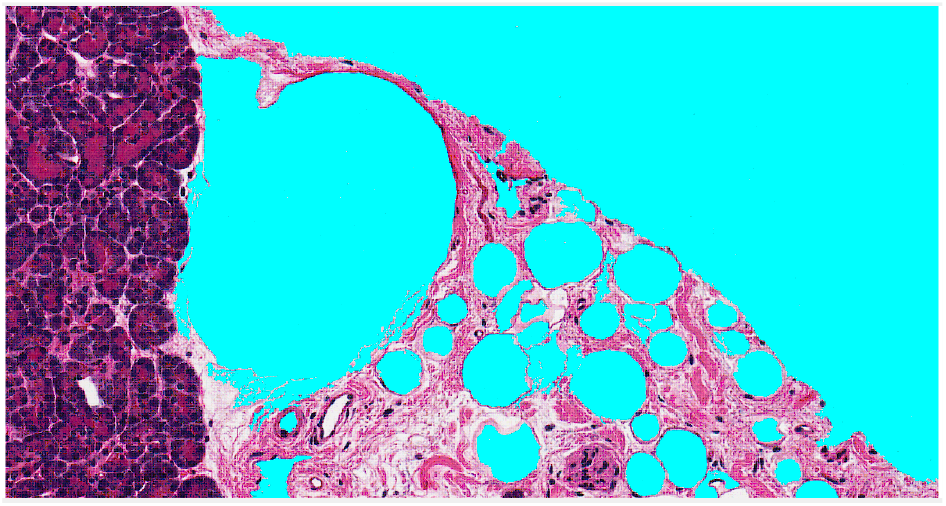

**Figure 13 (b).**
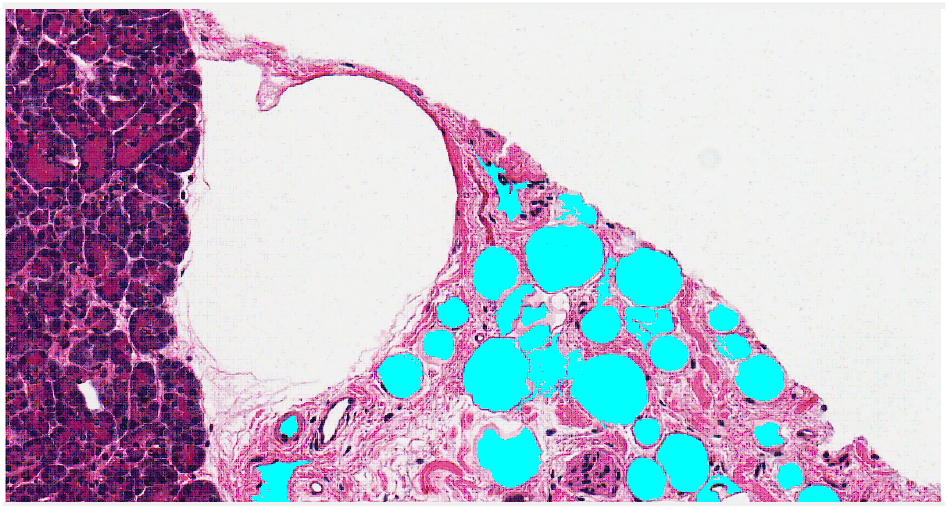

**Figure 14:**
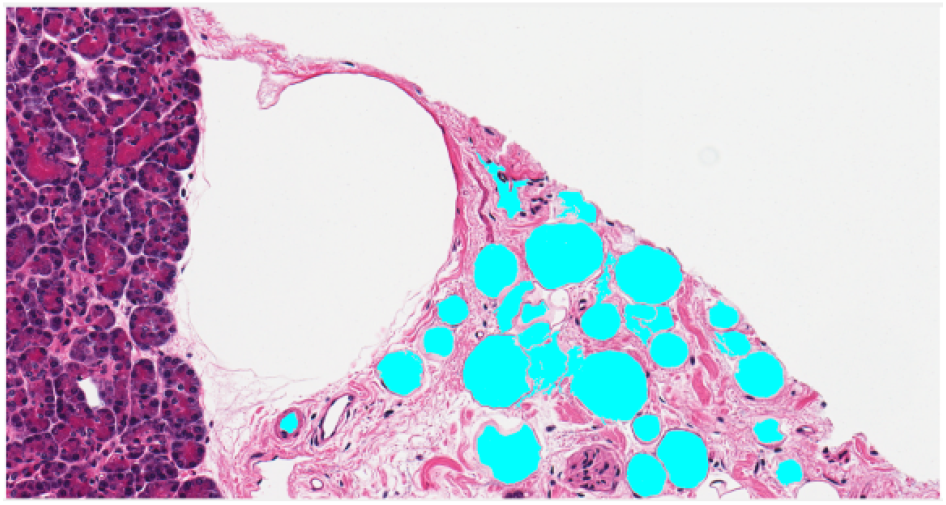
Fat cell removed from peripheral boundary.

#### 2.2.4. Removal of fats from boundary

Segments which are identified as fats but also contain pixels from boundary are discarded because their entire size is not known within the dimension of input image (Figure 14). While comparing Figure 14 with Figure 13 (b), it can be seen that some segments which contain pixels from bottom boundary gets discarded.

Some of these segments may even get discarded while selecting segments as per diameter. E.g. in Figure 13 (b), one big segment which has pixels in boundary gets discarded after considering maximum diameter area, whereas the segment is present in Figure 13 (a).

Removing fats from boundaries is the final step for tagging fats using machine. If users do not have manual tagged data of fat cells then they can conclude the experiment after this step.

#### 2.2.5. Analysis of valid fats

The machine tagged fat segments are compared with manually tagged fat segments to check the validity of our image analysis algorithm. If machine and manually tagged segments have pixels in common then, those pixels are considered as valid. The accuracy of the output is calculated in terms of Intersection over Union (IoU). The formula is:

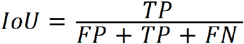

where TP = True Positive; FP = False Positive; FN = False Negative. TP is the intersection of machine and manually tagged pixels, FP are the pixels which are tagged by machine but do not become part of the intersection (i.e. Machine Tagged Area - TP) and FN are the pixels which are manually tagged but do not become part of the intersection (i.e. Manual Tagged Area - TP).

Figure 15 is the output generated after comparing machine and manual tagged fats. The Light Green colored pixels (RGB: 127, 255, 127) represents TP, Cyan color represents FN and Yellow color represents FP. Machine tagged, manual tagged and TP areas are 130,816, 111,300 and 110,429 pixels respectively. Hence as per the parameters used while demonstrating the steps, the IoU value is 0.838 but it can increase if parameters are changed or a better manual tagged image is referred.

**Figure 15.**
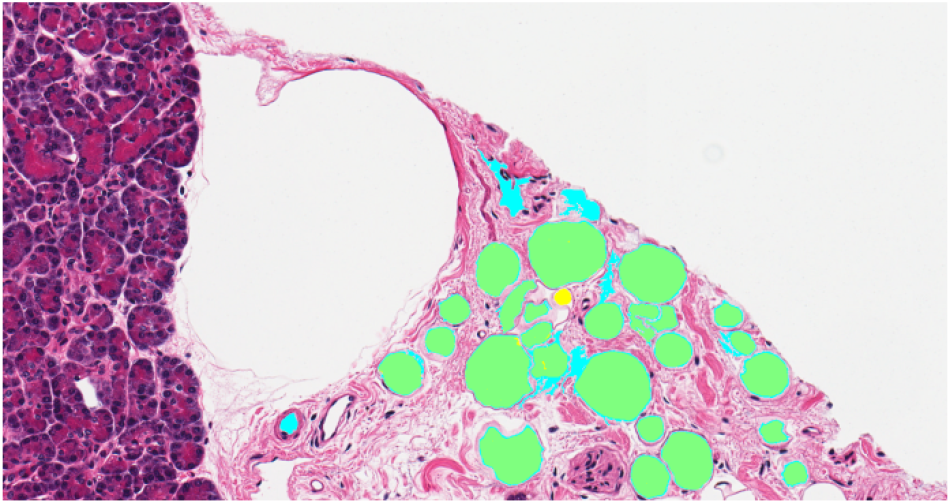

## 3. Results

Fat cell identification on ten sample images of dimension 1716 × 905 pixels was performed using Adiposoft and Fatquant tools. The outputs are shown below.

The threshold value, minimum and maximum diameter were chosen to get an optimal output.

Additionally, we analyzed liver images (Roy et al 2020) using our fatquant tool with IoU of 0.757 to 0.907 indicating wide applicability of our tool

Image (a) in Figures 16 – 25 are the raw sample images; (b) are manual fat tagged images; (c) and (d) are outputs from Adiposoft and Fatquant respectively. The parameter values of Adiposoft and Fatquant (mentioned in Table 1) are not exactly same because the default threshold or edge detection values used by Adiposoft are unknown to users. Hence for analysis, values have been chosen as per optimal output. Pixels being part of fat in (b) are marked with yellow color whereas pixels identified as fat in (c) and (d) are marked with cyan color.

**Figure 16.**
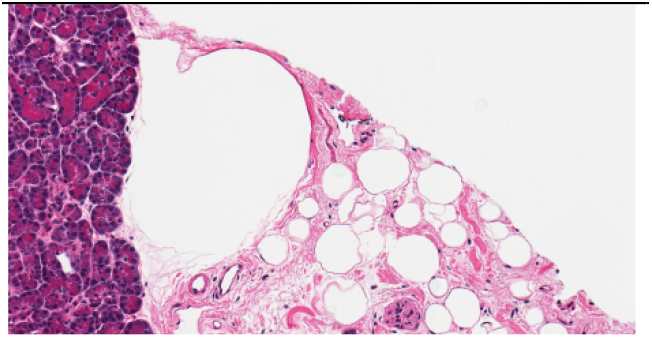
(a) Raw image.

**Figure 16.**
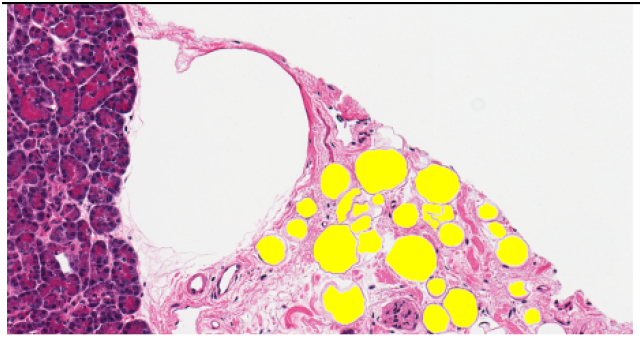
(b) Tagged image.

**Figure 16.**
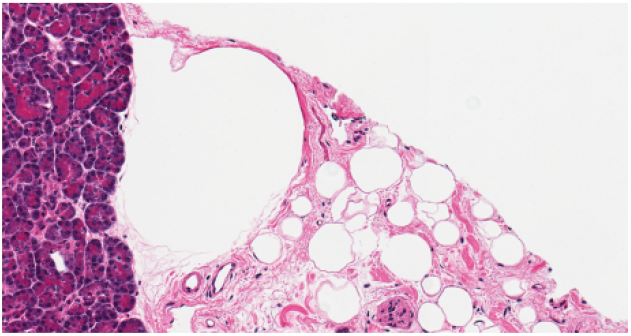
(c) Adiposoft.

**Figure 16.**
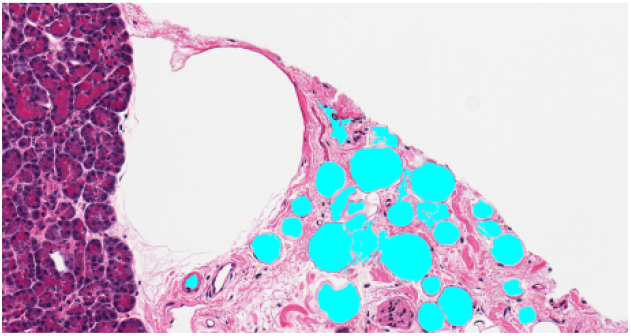
(d) Fatquant.

**Figure 17.**
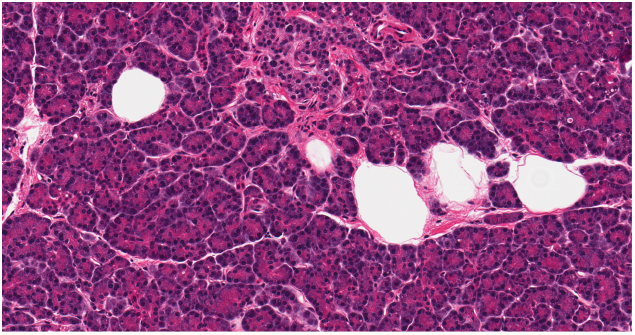
(a) Raw image.

**Figure 17.**
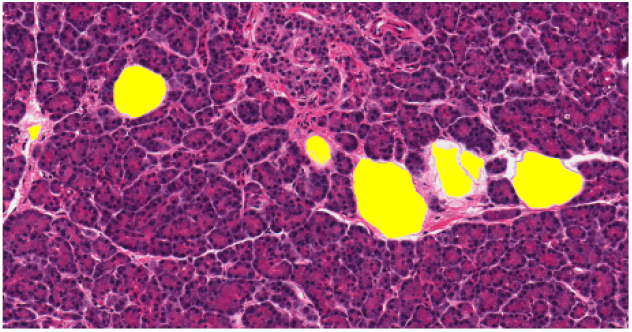
(b) Tagged image.

**Figure 17.**
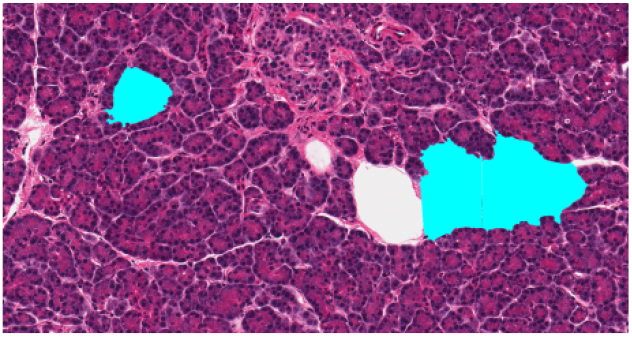
(c) Adiposoft.

**Figure 17.**
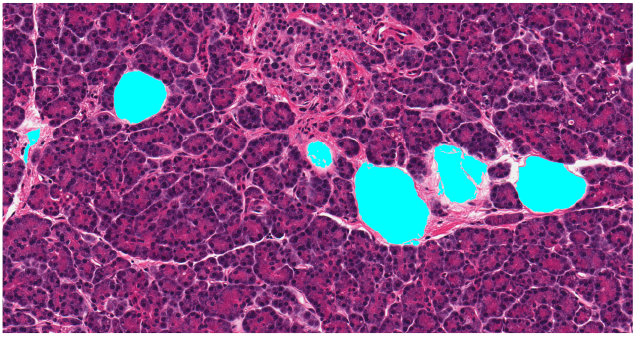
(d) Fatquant.

**Figure 18.**
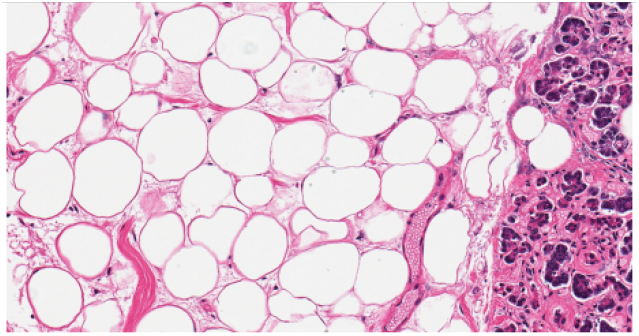
(a) Raw image.

**Figure 18.**
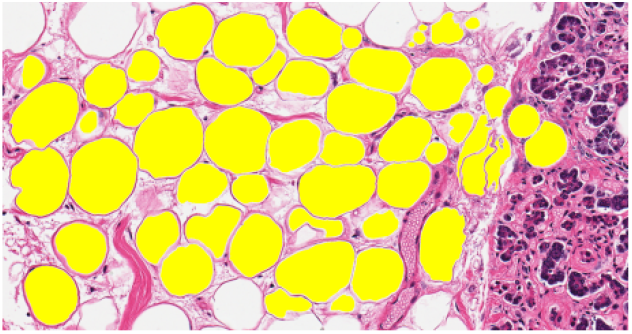
(b) Tagged image.

**Figure 18.**
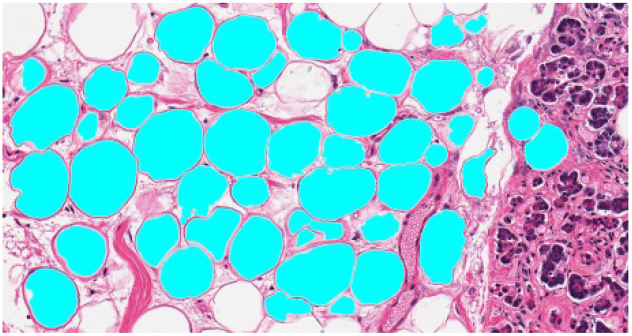
(c) Adiposoft.

**Figure 18.**
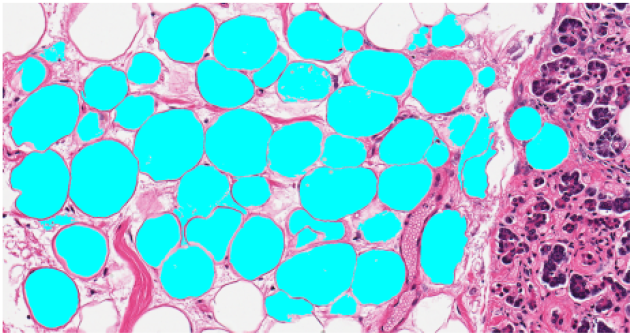
(d) Fatquant.

**Figure 19.**
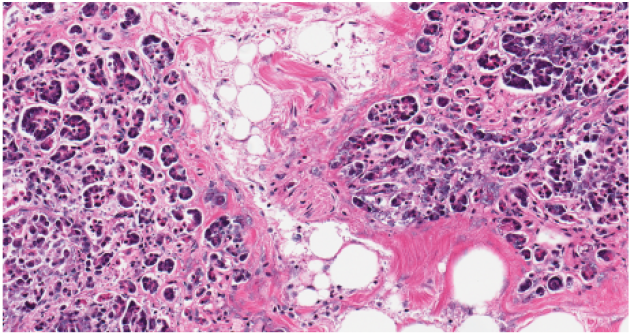
(a) Raw image.

**Figure 19.**
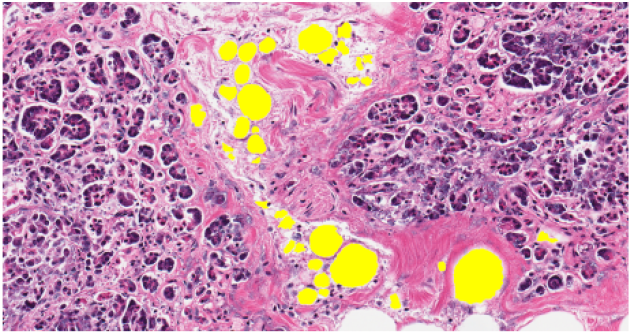
(b) Tagged image.

**Figure 19.**
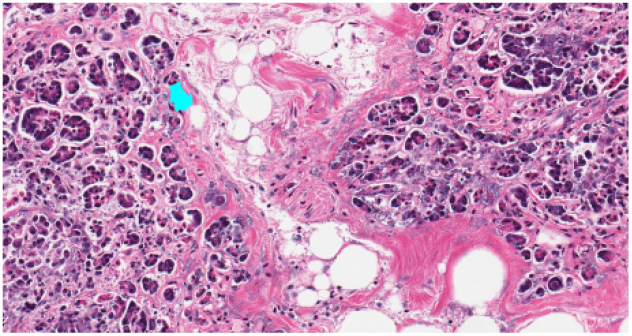
(c) Adiposoft.

**Figure 19.**
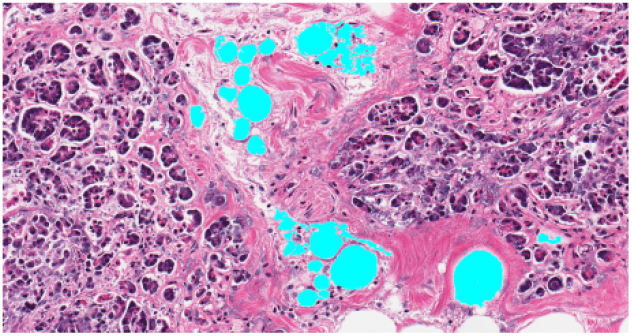
(d) Fatquant.

**Figure 20.**
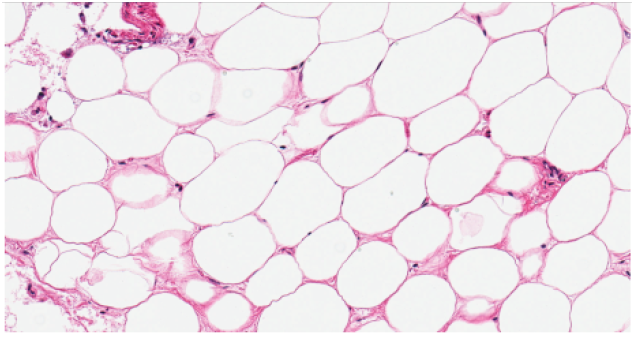
(a) Raw image.

**Figure 20.**
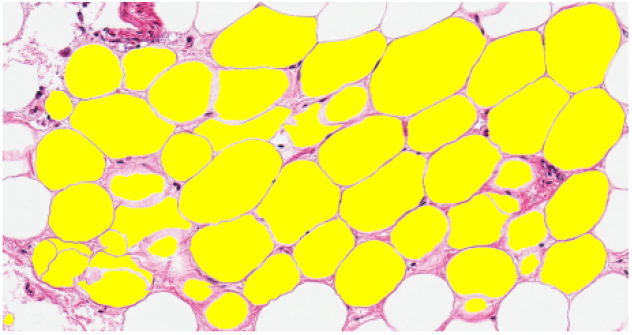
(b) Tagged image.

**Figure 20.**
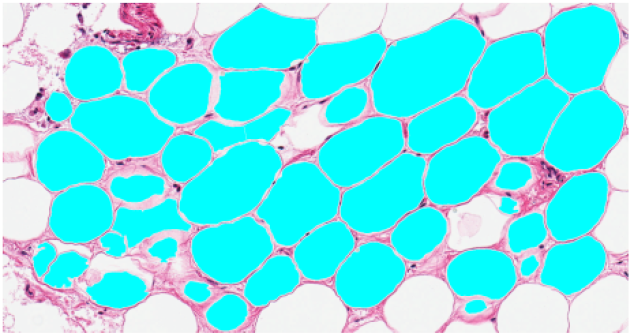
(c) Adiposoft.

**Figure 20.**
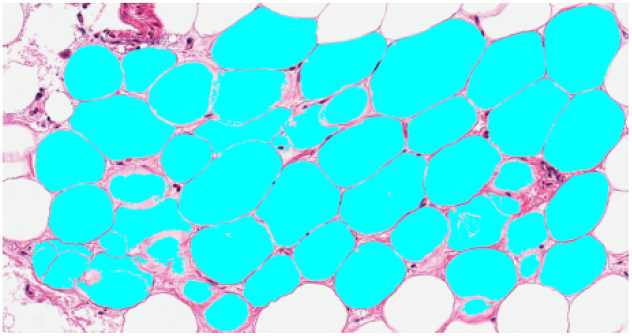
(d) Fatquant.

**Figure 21.**
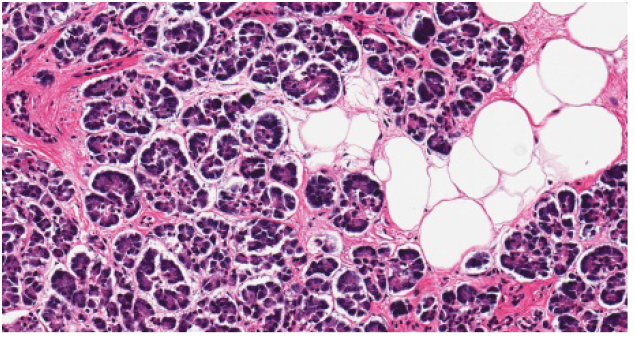
(a) Raw image.

**Figure 21.**
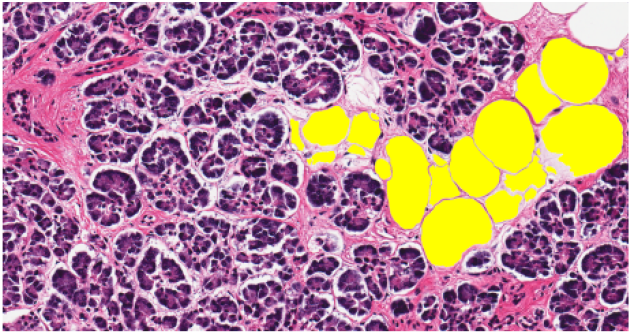
(b) Tagged image.

**Figure 21.**
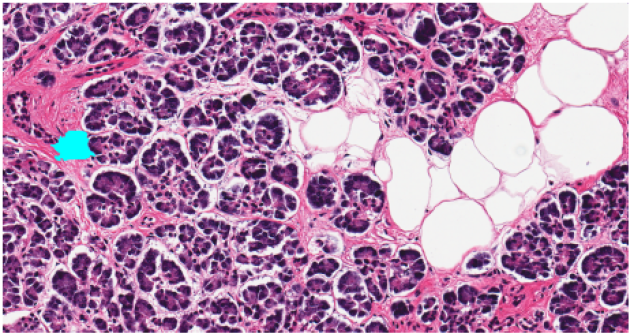
(c) Adiposoft.

**Figure 21.**
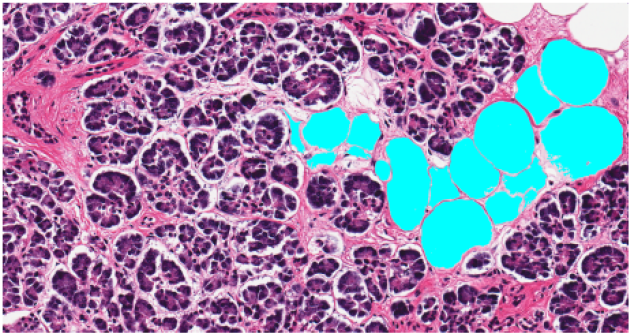
(d) Fatquant.

**Figure 22.**
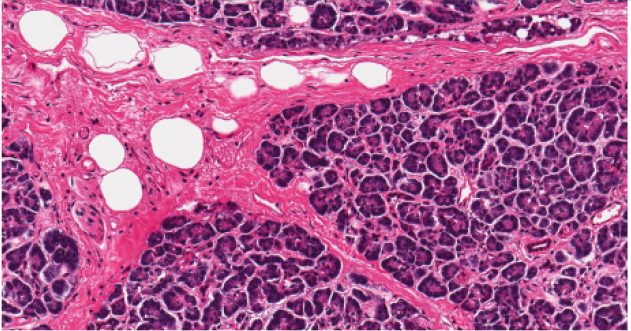
(a) Raw image.

**Figure 22.**
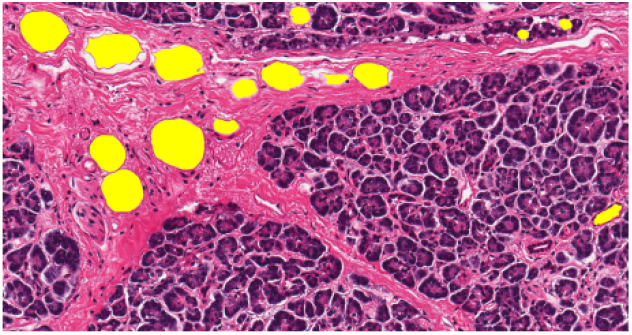
(b) Tagged image.

**Figure 22.**
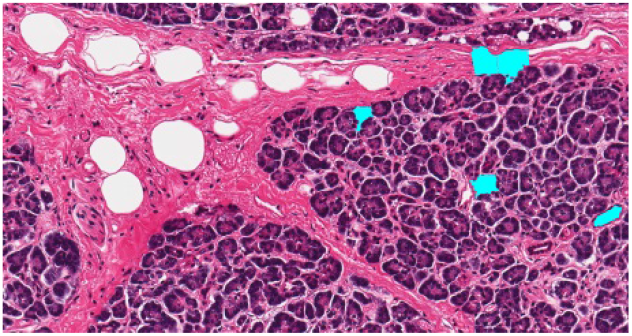
(c) Adiposoft.

**Figure 22.**
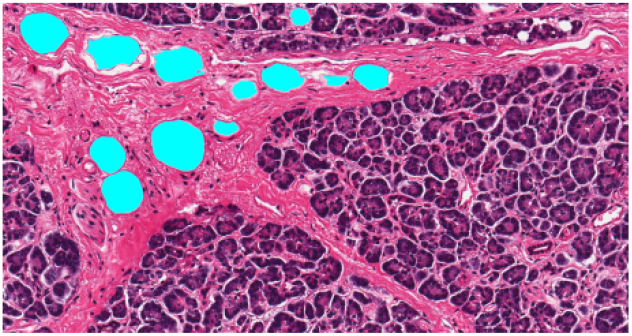
(d) Fatquant.

**Figure 23.**
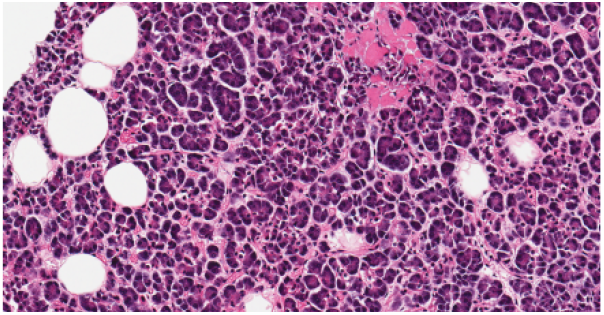
(a) Raw image.

**Figure 23.**
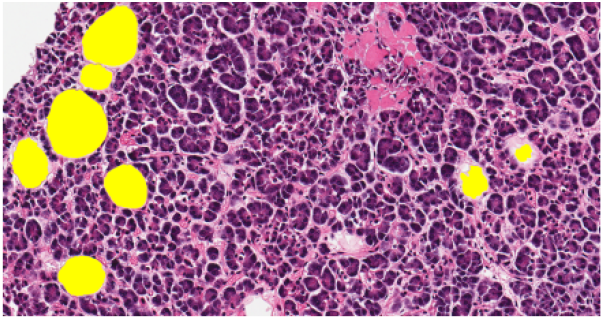
(b) Tagged image.

**Figure 23.**
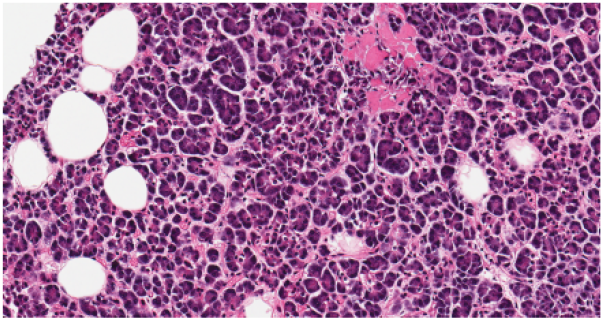
(c) Adiposoft.

**Figure 23.**
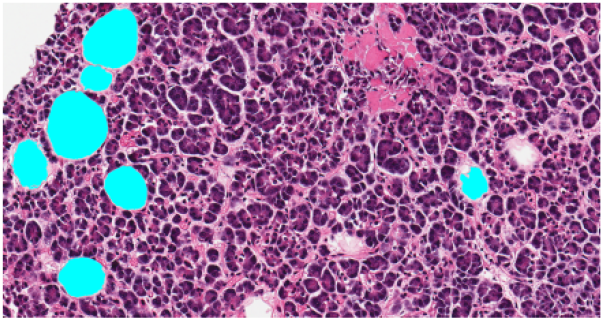
(d) Fatquant.

**Figure 24.**
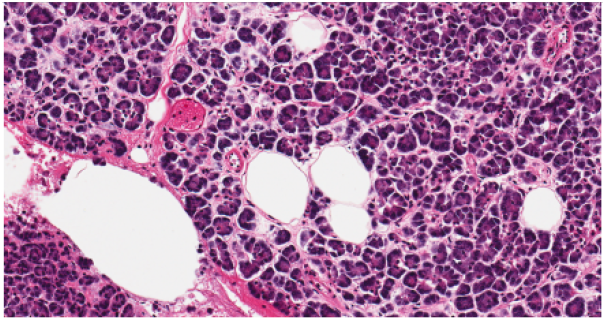
(a) Raw image.

**Figure 24.**
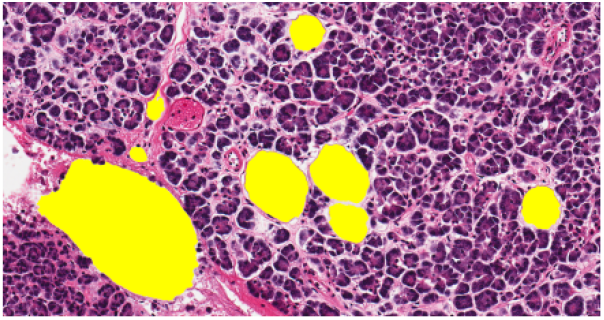
(b) Tagged image.

**Figure 24.**
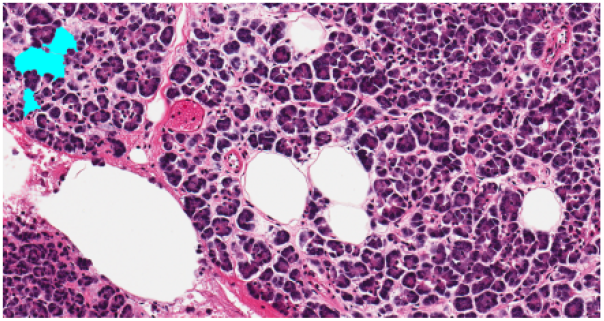
(c) Adiposoft.

**Figure 24.**
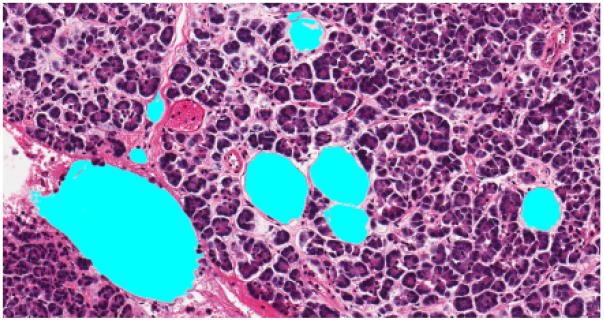
(d) Fatquant.

**Figure 25.**
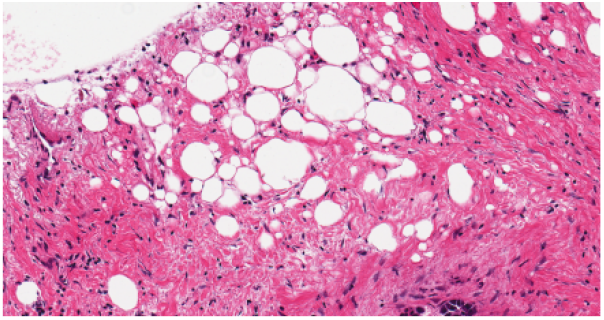
(a) Raw image.

**Figure 25.**
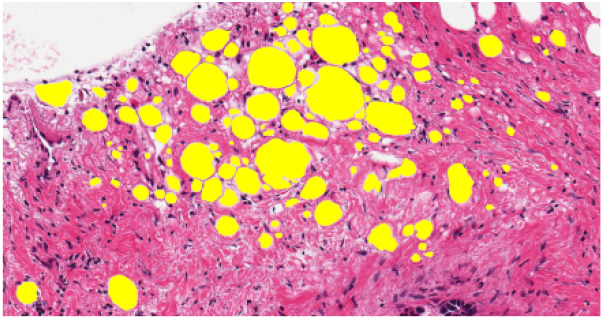
(b) Tagged image.

**Figure 25.**
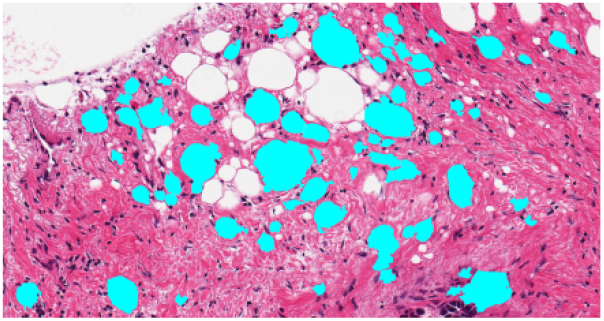
(c) Adiposoft.

**Figure 25.**
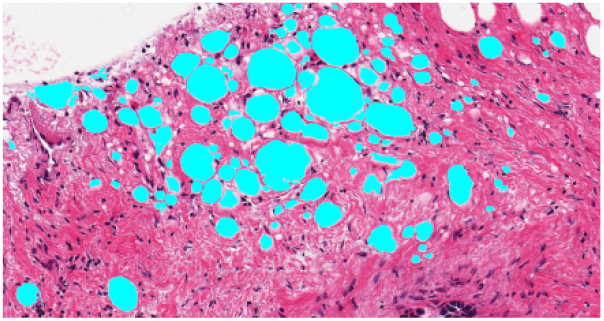
(d) Fatquant.

**Table 1:** Parameters used by tools in Figures 16 – 25

| Fig-<br>ures | Sam-<br>ples | GTEx<br>slide ID | Adiposoft |  |  | Fatquant |  |  |
| --- | --- | --- | --- | --- | --- | --- | --- | --- |
|  |  |  | Mi-<br>crons<br>per | Min.<br>diame-<br>ter | Max.<br>diame-<br>ter | Thresh-<br>old value | Min.<br>diameter | Max.<br>diameter |
|  |  |  | pixel |  |  |  |  |  |
| 16 | 1 | 11DXZ-0826 | 0.4942 | 20 | 135 | 230 | 27 | 130 |
| 17 | 2 | 1122O-0<br>726 | 0.4942 | 25 | 500 | 228 | 25 | 500 |
| 18 | 3 | 1117F-1<br>726 | 0.4942 | 22 | 500 | 228 | 29 | 500 |
| 19 | 4 | 1117F-1<br>726 | 0.4942 | 20 | 180 | 233 | 24 | 150 |
| 20 | 5 | 117YW-0926 | 0.4942 | 20 | 280 | 231 | 24 | 250 |
| 21 | 6 | 117YW-0926 | 0.4942 | 22 | 280 | 229 | 27 | 250 |
| 22 | 7 | 13PVQ-2026 | 0.4942 | 22 | 250 | 226 | 29 | 200 |
| 23 | 8 | 13FHP-1<br>926 | 0.4942 | 40 | 300 | 228 | 40 | 250 |
| 24 | 9 | 13FHP-1<br>926 | 0.4942 | 22 | 550 | 228 | 22 | 500 |
| 25 | 10 | 11WQC-0926 | 0.4942 | 14 | 200 | 231 | 14 | 200 |

**Table 2:**
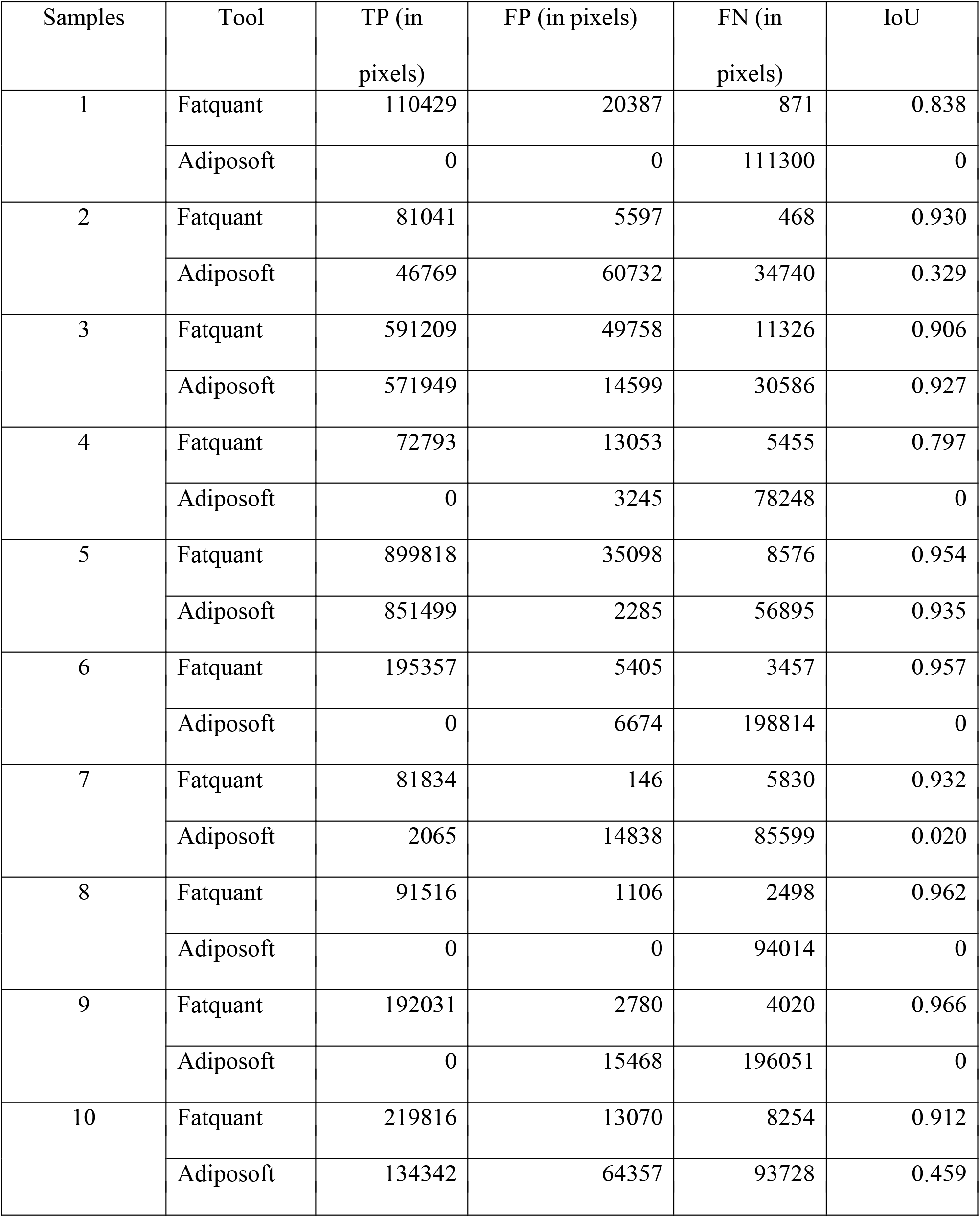
Calculation of accuracy on pancreas sample images

**Table 3:** Fatquant parameters for liver sample images

| Samples | Threshold value | Min. diameter | Max. diameter |
| --- | --- | --- | --- |
| 000 | 205 | 50 | 150 |
| 010 | 205 | 28 | 220 |
| 036 | 211 | 34 | 150 |
| 058 | 210 | 45 | 200 |
| 090 | 207 | 40 | 140 |
| 116 | 207 | 45 | 200 |
| 120 | 207 | 40 | 200 |
| 196 | 216 | 40 | 200 |
| 212 | 203 | 35 | 300 |
| 319 | 212 | 32 | 100 |
| 335 | 220 | 31 | 150 |
| 385 | 218 | 23 | 220 |
| 421 | 215 | 20 | 130 |
| 440 | 215 | 30 | 150 |
| 498 | 208 | 38 | 180 |
| 515 | 212 | 34 | 220 |
| 555 | 215 | 42 | 220 |
| 596 | 215 | 36 | 180 |
| 610 | 213 | 22 | 180 |
| 723 | 212 | 32 | 150 |

The threshold value, minimum and maximum diameter were chosen to get an optimal output.

**Table 4:** Calculation of accuracy using Fatquant on liver sample images

| Samples | TP (in pixels) | FP (in pixels) | FN (in pixels) | IoU |
| --- | --- | --- | --- | --- |
| 000 | 27496 | 5223 | 97 | 0.838 |
| 010 | 66999 | 10834 | 2419 | 0.835 |
| 036 | 21561 | 3940 | 129 | 0.841 |
| 058 | 39343 | 5233 | 91 | 0.881 |
| 090 | 61281 | 8386 | 510 | 0.873 |
| 116 | 50354 | 9816 | 607 | 0.828 |
| 120 | 65042 | 10587 | 94 | 0.859 |
| 196 | 119188 | 9727 | 2547 | 0.907 |
| 212 | 62885 | 12752 | 298 | 0.828 |
| 319 | 20594 | 6103 | 511 | 0.757 |
| 335 | 45856 | 10862 | 1614 | 0.786 |
| 385 | 56330 | 9776 | 1268 | 0.836 |
| 421 | 73060 | 12052 | 326 | 0.855 |
| 440 | 28788 | 4726 | 677 | 0.842 |
| 498 | 63565 | 12067 | 1301 | 0.826 |
| 515 | 96819 | 12566 | 1514 | 0.873 |
| 555 | 69386 | 12856 | 1222 | 0.831 |
| 596 | 21020 | 4502 | 528 | 0.807 |
| 610 | 52206 | 10069 | 1852 | 0.814 |
| 723 | 12971 | 3286 | 436 | 0.777 |

From the outputs it can be noted that Adiposoft only shows decent output when adipocytes cover maximum area of a sample image (e.g. Figures 18 and 20). In a heterogeneous sample image this tool can tag many non-fat areas as valid fats (e.g. Figures 21, 22 and 24). Moreover, it can even fail to identify presence of any fat in an image (e.g. Figures 16 and 23). Hence, Fatquant performs better in the scenarios shown. The tagging of fat cells shown in (b) images are done by the authors and not collected from any laboratory. Ground truth data of the slides used was not officially made available by the source.

Image (a) in Figures 26 – 31 are the sample images; (b) represents manual tagged and (c) represents machine tagged areas using Fatquant of images in (a). As per the data mentioned in Table 5 it can be noted that fat cells available in Figures 26 – 28 are easy to get tagged by this tool, hence IoU value increases. Whereas the tool does not detect fat cell properly when cell boundaries are not clearly delineated as seen in Figures 29 – 31, hence IoU value decreases. So validity analysis performed on sample images with many fat cells similar to that of Figures 26 – 28 will likely show higher accuracy. Manual tagged data shown in (b) images are not collected from any laboratory website and are rather created by the authors. So, there can be some variation in ground truth data of these images created by any other source. In the GitHub repository mentioned, sample images used in Figures 16 – 25 and Figures 26 – 31 are available in ‘Test_samples’ and ‘Small_samples’ directories respectively.

**Figure 26.**
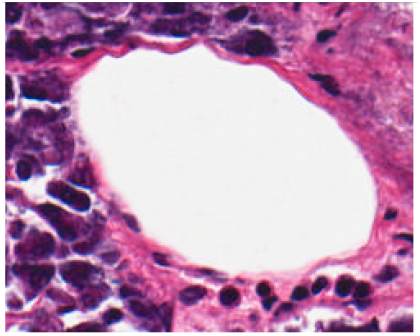
(a) sample.

**Figure 26.**
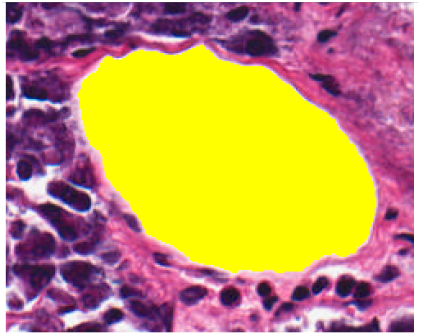
(b) tagged.

**Figure 26.**
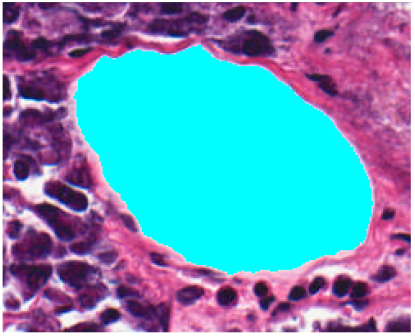
(c) Fatquant.

**Figure 27.**
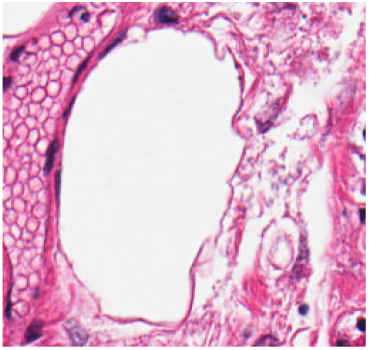
(a) sample.

**Figure 27.**
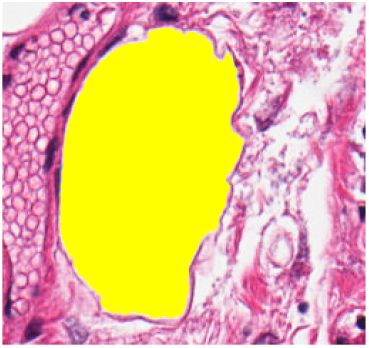
(b) tagged.

**Figure 27.**
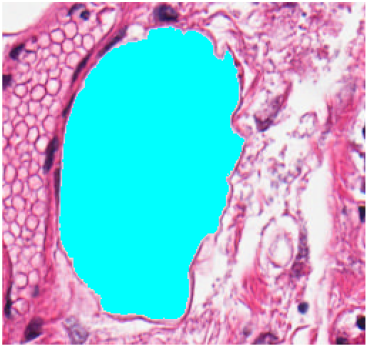
(c) Fatquant.

**Figure 28.**
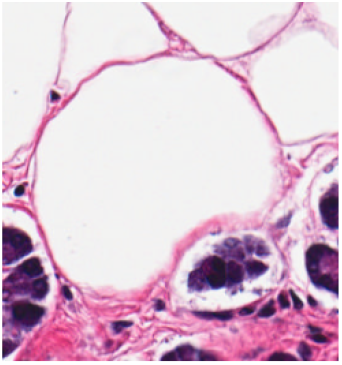
(a) sample.

**Figure 28.**
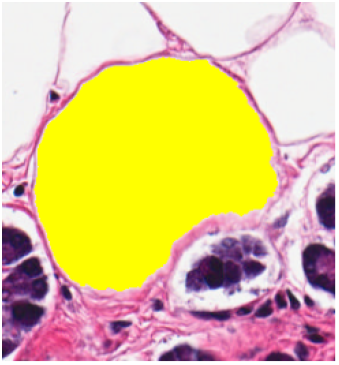
(b) tagged.

**Figure 28.**
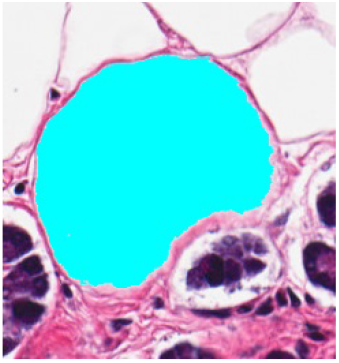
(c) Fatquant.

**Figure 29.**
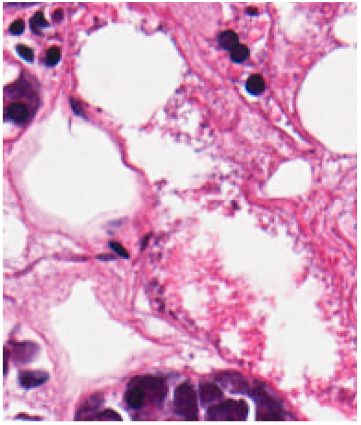
(a) sample.

**Figure 29.**
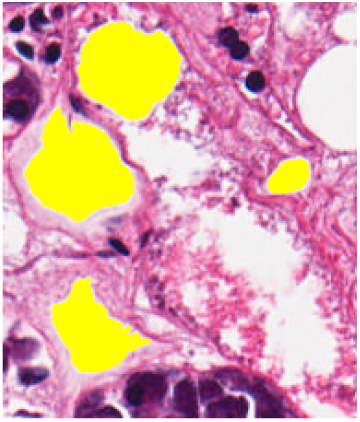
(b) tagged.

**Figure 29.**
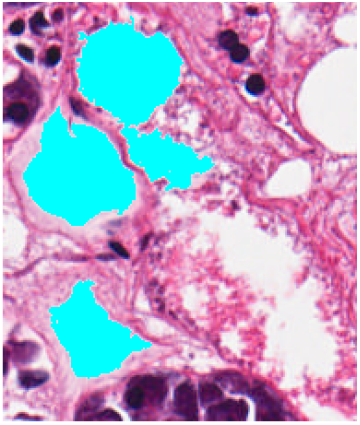
(c) Fatquant.

**Figure 30.**
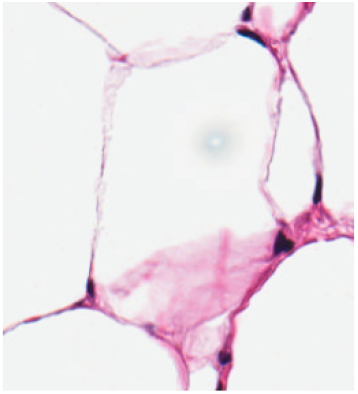
(a) sample.

**Figure 30.**
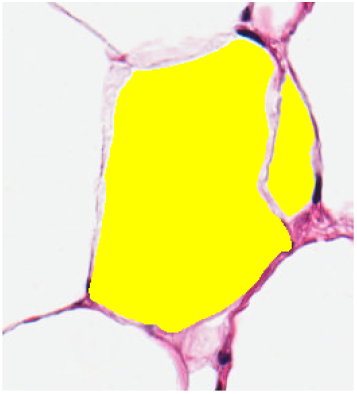
(b) tagged.

**Figure 30.**
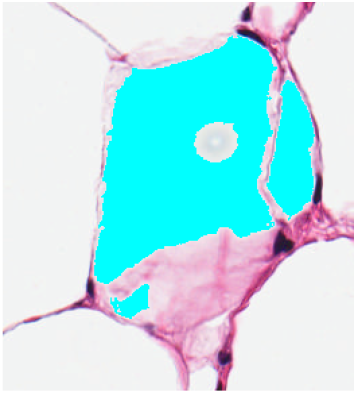
(c) Fatquant.

**Figure 31.**
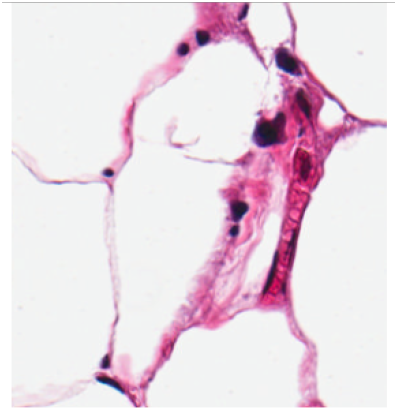
(a) sample.

**Figure 31.**
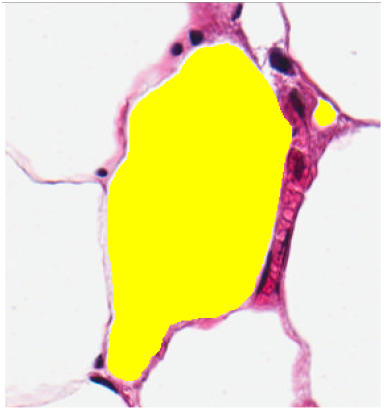
(b) tagged.

**Figure 31.**
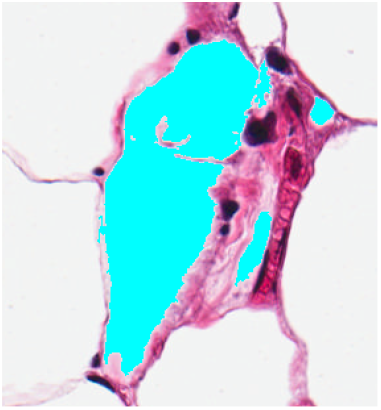
(c) Fatquant.

**Table 5:** Accuracy of machine annotation by Fatquant in Figures 26 – 31

| Fig-<br>ures | GTEx<br>slide ID | Thresh-<br>old value | Min.<br>diameter | Max.<br>diameter | TP (in<br>pixels) | FP (in<br>pixels) | FN (in<br>pixels) | IoU |
| --- | --- | --- | --- | --- | --- | --- | --- | --- |
| 26 | 117YW<br>-0926 | 229 | 20 | 180 | 17640 | 118 | 102 | 0.987 |
| 27 | 1117F-1 | 227 | 30 | 200 | 19321 | 269 | 15 | 0.985 |
|  | 726 |  |  |  |  |  |  |  |
| 28 | 117YW<br>-0926 | 228 | 20 | 200 | 26095 | 115 | 334 | 0.983 |
| 29 | 111VG-0926 | 227 | 35 | 100 | 5692 | 1312 | 326 | 0.776 |
| 30 | 117YW<br>-0926 | 236 | 15 | 220 | 18178 | 499 | 7042 | 0.707 |
| 31 | 117YW<br>-0926 | 231 | 10 | 200 | 16441 | 484 | 7623 | 0.670 |

## 4. Discussion

Pancreatic fat accumulation has been associated with obesity, impaired b-cell function and may be an early sign in the development of metabolic syndrome (Dite et al 2020, Sequeira et al 2022, Rugivarodom et al 2022). Pancreas is a heterogenous tissue and current tools for fat cell analysis such as Adiposoft (Galarraga et al 2012), AdipoCount (Zhi et al 2018) are unable to analyze fat cell infiltration in pancreas. Current automated pancreas tool are largely restricted to segmentation of the endocrine islet fractions of pancreas and are based on cell nulcei identification, extracting colour features mostly for pancreatic cancer detection (Huang et al 2016, Vu et al 2019, Yang et al 2021). There are other deep learning based methods for pancreas segmentation but again this are largely restricted to cancer diagnosis (Huang et al 2021) but cannot analyze fatty cell infiltration.

We have developed a tool, Fatquant, to identify fat cells in heterogeneous histological tissue sections that can complement work of pathologist in identification and analysis of fat cells due to their growing importance in pathophysiological functions and with the availability of whole slide digitized images it can save significant amount of time. Based on our analysis, Adiposoft shows decent output when adipocytes cover maximum area of a sample image and label non-fat areas as valid fat cells in heterogenous tissue sample images. In absence of ground truth images, we manually annotated fat cells in different pancreas sample images and then compared the output accuracy using IoU with Fatquant tool annotated results. We observed an IoU of 0.797 to 0.966 exhibiting high degree of similarity between manual versus Fatquant annotation. Our fatquant tool works well for both homogenous adipose tissue and varied heterogenous tissues like pancreas, liver suggesting diverse applicability of the tool for analysis of fat cells.

## Acknowledgments

There is no potential conflicts of interest and no funding to declare. This research did not receive any specific grant from funding agencies in the public, commercial, or not-for-profit sectors.

